## Supplemental Table 1 for "Sphingosine-1-phosphate signaling regulates the ability of Müller glia to become neurogenic, proliferating progenitor-like cells"

**Supplemental table – Statistics for dotplots**

**Figure 1j - stats MG aggregate for S1P-related genes**

FindMarkers(NEW_MG_AGG, ident.1 = "resting MG", ident.2 = "24hr activated MG", features = c("S1PR1", "S1PR3", "SPHK1", "SGPL1", "ASAH1", "CERS5", "CERS6"), logfc.threshold = 0.05, min.pct = 0.01, test.use = "wilcox")

|++++++++++++++++++++++++++++++++++++++++++++++++++| 100% elapsed=00s

p_val avg_log2FC pct.1 pct.2	p_val_adj

S1PR1 0.000000e+00 1.4766129 0.547 0.224 0.000000e+00

CERS6 6.663376e-149 0.7696331 0.284 0.151 1.547569e-144

CERS5 4.614283e-70 -0.1341384 0.053 0.148 1.071667e-65

S1PR3 5.389332e-38 -0.0665710 0.026 0.076 1.251672e-33

SPHK1 1.423129e-06 0.2631905 0.120 0.105 3.305217e-02

SGPL1 2.876968e-04 0.2414398 0.116 0.106 1.000000e+00

FindMarkers(NEW_MG_AGG, ident.1 = "resting MG", ident.2 = c("MGPC 1", "MGPC 2", "MGPC 3"), features = c("S1PR1", "S1PR3", "SPHK1", "SGPL1", "ASAH1", "CERS5", "CERS6"), logfc.threshold = 0.05, min.pct = 0.01, test.use = "wilcox")

|++++++++++++++++++++++++++++++++++++++++++++++++++| 100% elapsed=00s

p_val avg_log2FC pct.1 pct.2	p_val_adj

S1PR1 0.000000e+00 1.27051799 0.547 0.241 0.000000e+00

CERS6 5.102892e-107 0.61536607 0.284 0.176 1.185147e-102

S1PR3 9.919697e-54 -0.14486671 0.026 0.087 2.303850e-49

CERS5 5.006465e-29 -0.07639374 0.053 0.103 1.162752e-24

SPHK1 6.348267e-21 0.25778520 0.120 0.085 1.474385e-16

SGPL1 8.274883e-01 0.10536822 0.116 0.122 1.000000e+00

FindMarkers(NEW_MG_AGG, ident.1 = c("return to resting MG", "returning farther toward resting MG"), ident.2 = c("MGPC 1", "MGPC 2", "MGPC 3"), features = c("S1PR1", "S1PR3", "SPHK1", "SGPL1", "ASAH1", "CERS5", "CERS6"), logfc.threshold = 0.05, min.pct = 0.01, test.use = "wilcox")

|++++++++++++++++++++++++++++++++++++++++++++++++++| 100% elapsed=00s

p_val avg_log2FC pct.1 pct.2	p_val_adj

CERS5 1.513637e-138 -0.18156704 0.030 0.103 3.515422e-134

S1PR1 3.034636e-65 0.43425708 0.294 0.241 7.047942e-61

SGPL1 8.094624e-02 0.08006903 0.111 0.122 1.000000e+00

CERS6 2.902173e-01 0.15002088 0.160 0.176 1.000000e+00

FindMarkers(NEW_MG_AGG, ident.1 = c("2do FI"), ident.2 = c("24hr activated MG"), features = c("S1PR1", "S1PR3", "SPHK1", "SGPL1", "ASAH1", "CERS5", "CERS6"), logfc.threshold = 0.05, min.pct = 0.01, test.use = "wilcox")

|++++++++++++++++++++++++++++++++++++++++++++++++++| 100% elapsed=00s

p_val avg_log2FC pct.1 pct.2	p_val_adj

ASAH1 5.786758e-197 -0.3543778 0.149 0.368 1.343974e-192

S1PR1 1.685156e-157 -0.2671338 0.071 0.224 3.913775e-153

SPHK1 3.115854e-84 -0.1404222 0.029 0.105 7.236571e-80

S1PR3 3.117572e-81 -0.1468519 0.013 0.076 7.240562e-77

CERS5 1.993400e-80 -0.1385227 0.056 0.148 4.629670e-76

CERS6 1.588703e-76 -0.1281614 0.060 0.151 3.689762e-72

SGPL1 5.346322e-01 0.1722083 0.103 0.106 1.000000e+00

FindMarkers(NEW_MG_AGG, ident.1 = c("mix activated MG"), ident.2 = c("MGPC 1", "MGPC 2", "MGPC 3"), features = c("S1PR1", "S1PR3", "SPHK1", "SGPL1", "ASAH1", "CERS5", "CERS6"), logfc.threshold = 0.05, min.pct = 0.01, test.use = "wilcox")

|++++++++++++++++++++++++++++++++++++++++++++++++++| 100% elapsed=00s

p_val avg_log2FC pct.1 pct.2	p_val_adj

CERS5 7.958113e-55 -0.07274424 0.058 0.103 1.848272e-50

S1PR3 6.163136e-49 -0.07884567 0.048 0.087 1.431388e-44

CERS6 3.410463e-36 -0.08659888 0.127 0.176 7.920801e-32

SGPL1 8.008423e-03 0.05147612 0.112 0.122 1.000000e+00

S1PR1 4.978803e-01 0.09742839 0.230 0.241 1.000000e+00

**Figure 1m - stats MG early time points for S1P-related genes**

FindMarkers(early_MG, ident.1 = "saline", ident.2 = "NMDA 3hr", group.by = "orig.ident", features = c("S1PR1", "ASAH1", "SGPL1", "SGPP1", "CERS5", "CERS6"), min.pct = 0.001, logfc.threshold = 0.001, test.use = "wilcox")

|++++++++++++++++++++++++++++++++++++++++++++++++++| 100% elapsed=00s

p_val avg_log2FC pct.1 pct.2	p_val_adj

S1PR1 4.861920e-74 1.35840824 0.594 0.264 9.061647e-70

CERS6 2.927224e-50 -0.50048518 0.319 0.638 5.455761e-46

SGPP1 4.748836e-05 0.01775621 0.089 0.149 8.850881e-01

SGPL1 6.107935e-01 0.16573900 0.273 0.284 1.000000e+00

CERS5 6.721510e-01 0.04612966 0.122 0.133 1.000000e+00

ASAH1 7.408299e-01 0.22668713 0.526 0.601 1.000000e+00

FindMarkers(early_MG, ident.1 = "saline", ident.2 = "NMDA 12hr", group.by = "orig.ident", features = c("S1PR1", "ASAH1", "SGPL1", "SGPP1", "CERS5", "CERS6"), min.pct = 0.001, logfc.threshold = 0.001, test.use = "wilcox")

|++++++++++++++++++++++++++++++++++++++++++++++++++| 100% elapsed=00s

p_val avg_log2FC pct.1 pct.2	p_val_adj

S1PR1 2.075700e-182 1.56812286 0.594 0.145 3.868690e-178

CERS6 4.553748e-19 0.29797231 0.319 0.195 8.487276e-15

ASAH1 1.779123e-07 0.51559364 0.526 0.534 3.315930e-03

CERS5 5.699382e-05 -0.04947385 0.122 0.175 1.000000e+00

SGPP1 1.899173e-02 0.06397734 0.089 0.115 1.000000e+00

SGPL1 9.009138e-01 0.14016846 0.273 0.286 1.000000e+00

FindMarkers(early_MG, ident.1 = "saline", ident.2 = "NMDA 48hr", group.by = "orig.ident", features = c("S1PR1", "ASAH1", "SGPL1", "SGPP1", "CERS5", "CERS6"), min.pct = 0.001, logfc.threshold = 0.001, test.use = "wilcox")

|++++++++++++++++++++++++++++++++++++++++++++++++++| 100% elapsed=00s

p_val avg_log2FC pct.1 pct.2	p_val_adj

S1PR1 1.084987e-52 0.96089885 0.594 0.382 2.022199e-48

SGPP1 5.052564e-07 0.01096686 0.089 0.161 9.416968e-03

CERS5 4.636410e-06 -0.01106706 0.122 0.202 8.641341e-02

SGPL1 3.410185e-03 -0.01281733 0.273 0.356 1.000000e+00

CERS6 8.645303e-02 0.09526313 0.319 0.325 1.000000e+00

ASAH1 2.039700e-01 0.11915563 0.526 0.616 1.000000e+00

FindMarkers(early_MG, ident.1 = "NMDA 3hr", ident.2 = "NMDA 48hr", group.by = "orig.ident", features = c("S1PR1", "ASAH1", "SGPL1", "SGPP1", "CERS5", "CERS6"), min.pct = 0.001, logfc.threshold = 0.001, test.use = "wilcox")

|++++++++++++++++++++++++++++++++++++++++++++++++++| 100% elapsed=00s

p_val	avg_log2FC pct.1 pct.2	p_val_adj

CERS6 5.321084e-53 0.595748308 0.638 0.325 9.917437e-49

S1PR1 4.956422e-07 -0.397509391 0.264 0.382 9.237779e-03

CERS5 1.655711e-03 -0.057196716 0.133 0.202 1.000000e+00

SGPL1 6.708620e-03 -0.178556331 0.284 0.356 1.000000e+00

ASAH1 1.202453e-01 -0.107531503 0.601 0.616 1.000000e+00

SGPP1 7.985372e-01 -0.006789352 0.149 0.161 1.000000e+00

FindMarkers(early_MG, ident.1 = "NMDA 12hr", ident.2 = "NMDA 48hr", group.by = "orig.ident", features = c("S1PR1", "ASAH1", "SGPL1", "SGPP1", "CERS5", "CERS6"), min.pct = 0.001, logfc.threshold = 0.001, test.use = "wilcox")

|++++++++++++++++++++++++++++++++++++++++++++++++++| 100% elapsed=00s

p_val avg_log2FC pct.1 pct.2	p_val_adj

S1PR1 2.913074e-41 -0.60722401 0.145 0.382 5.429387e-37

ASAH1 1.112567e-10 -0.39643801 0.534 0.616 2.073603e-06

CERS6 1.726546e-10 -0.20270918 0.195 0.325 3.217937e-06

SGPL1 4.570531e-03 -0.15298579 0.286 0.356 1.000000e+00

SGPP1 7.408533e-03 -0.05301048 0.115 0.161 1.000000e+00

CERS5 4.896419e-01 0.03840679 0.175 0.202 1.000000e+00

FindMarkers(early_MG, ident.1 = "NMDA 3hr", ident.2 = "NMDA 12hr", group.by = "orig.ident", features = c("S1PR1", "ASAH1", "SGPL1", "SGPP1", "CERS5", "CERS6"), min.pct = 0.001, logfc.threshold = 0.001, test.use = "wilcox")

|++++++++++++++++++++++++++++++++++++++++++++++++++| 100% elapsed=00s

p_val avg_log2FC pct.1 pct.2	p_val_adj

CERS6 1.016448e-106 0.79845749 0.638 0.195 1.894456e-102

S1PR1 1.276494e-11 0.20971462 0.264 0.145 2.379129e-07

ASAH1 6.926298e-05 0.28890651 0.601 0.534 1.000000e+00

CERS5 7.040153e-03 -0.09560351 0.133 0.175 1.000000e+00

SGPP1 4.137025e-02 0.04622113 0.149 0.115 1.000000e+00

SGPL1 6.688715e-01 -0.02557054 0.284 0.286 1.000000e+00

**Figure 1n - stats retinal cells early time points for S1P-related genes**

**Whole retina**

FindMarkers(early_AGG, ident.1 = "saline", ident.2 = "NMDA 3hr", group.by = "orig.ident", features = c("S1PR1", "ASAH1", "SGPL1", "SGPP1", "CERS5", "CERS6"), min.pct = 0.001, logfc.threshold = 0.001, test.use = "wilcox")

|++++++++++++++++++++++++++++++++++++++++++++++++++| 100% elapsed=00s

p_val avg_log2FC pct.1 pct.2	p_val_adj

S1PR1 3.997171e-74 0.47243542 0.128 0.044 7.449927e-70

CERS6 7.500436e-05 -0.06079080 0.136 0.159 1.000000e+00

SGPP1 1.643961e-03 -0.05882122 0.068 0.082 1.000000e+00

SGPL1 2.805797e-02 0.03611440 0.142 0.132 1.000000e+00

CERS5 2.733502e-01 -0.01319053 0.063 0.058 1.000000e+00

ASAH1 9.554875e-01 -0.08849468 0.266 0.265 1.000000e+00

FindMarkers(early_AGG, ident.1 = "saline", ident.2 = "NMDA 12hr", group.by = "orig.ident", features = c("S1PR1", "ASAH1", "SGPL1", "SGPP1", "CERS5", "CERS6"), min.pct = 0.001, logfc.threshold = 0.001, test.use = "wilcox")

|++++++++++++++++++++++++++++++++++++++++++++++++++| 100% elapsed=00s

p_val avg_log2FC pct.1 pct.2	p_val_adj

S1PR1 1.255749e-196 0.51656004 0.128 0.029 2.340464e-192

CERS6 3.061930e-44 0.17922635 0.136 0.081 5.706824e-40

ASAH1 5.239887e-06 0.23777841 0.266 0.248 9.766102e-02

SGPL1 5.362245e-04 0.01485817 0.142 0.127 1.000000e+00

CERS5 1.685029e-03 0.02389932 0.063 0.053 1.000000e+00

SGPP1 3.418975e-02 0.01707665 0.068 0.061 1.000000e+00

FindMarkers(early_AGG, ident.1 = "NMDA 3hr", ident.2 = "NMDA 12hr", group.by = "orig.ident", features = c("S1PR1", "ASAH1", "SGPL1", "SGPP1", "CERS5", "CERS6"), min.pct = 0.001, logfc.threshold = 0.001, test.use = "wilcox")

|++++++++++++++++++++++++++++++++++++++++++++++++++| 100% elapsed=00s

p_val avg_log2FC pct.1 pct.2	p_val_adj

CERS6 1.701158e-60 0.24001715 0.159 0.081 3.170619e-56

S1PR1 5.140161e-08 0.04412462 0.044 0.029 9.580233e-04

SGPP1 3.380974e-07 0.07589787 0.082 0.061 6.301459e-03

ASAH1 2.396886e-04 0.32627309 0.265 0.248 1.000000e+00

CERS5 1.396240e-01 0.03708985 0.058 0.053 1.000000e+00

SGPL1 5.157213e-01 -0.02125623 0.132 0.127 1.000000e+00

FindMarkers(early_AGG, ident.1 = "NMDA 3hr", ident.2 = "NMDA 48hr", group.by = "orig.ident", features = c("S1PR1", "ASAH1", "SGPL1", "SGPP1", "CERS5", "CERS6"), min.pct = 0.001, logfc.threshold = 0.001, test.use = "wilcox")

|++++++++++++++++++++++++++++++++++++++++++++++++++| 100% elapsed=00s

p_val avg_log2FC pct.1 pct.2	p_val_adj

CERS6 3.439922e-25 0.22179458 0.159 0.105 6.411326e-21

SGPL1 9.646975e-07 -0.09531840 0.132 0.159 1.798003e-02

S1PR1 2.013430e-06 -0.07335224 0.044 0.062 3.752630e-02

SGPP1 5.707874e-03 0.07957794 0.082 0.070 1.000000e+00

ASAH1 1.344348e-01 0.05351781 0.265 0.278 1.000000e+00

CERS5 5.456803e-01 0.04543650 0.058 0.061 1.000000e+00

FindMarkers(early_AGG, ident.1 = "NMDA 12hr", ident.2 = "NMDA 48hr", group.by = "orig.ident", features = c("S1PR1", "ASAH1", "SGPL1", "SGPP1", "CERS5", "CERS6"), min.pct = 0.001, logfc.threshold = 0.001, test.use = "wilcox")

|++++++++++++++++++++++++++++++++++++++++++++++++++| 100% elapsed=00s

p_val	avg_log2FC pct.1 pct.2	p_val_adj

S1PR1 5.955411e-34 -0.117476858 0.029 0.062 1.109969e-29

SGPL1 1.188265e-11 -0.074062172 0.127 0.159 2.214689e-07

ASAH1 5.021067e-10 -0.272755282 0.248 0.278 9.358264e-06

CERS6 3.174457e-09 -0.018222573 0.081 0.105 5.916554e-05

CERS5 1.406153e-02 0.008346652 0.053 0.061 1.000000e+00

SGPP1 1.418724e-02 0.003680066 0.061 0.070 1.000000e+00

**Microglia**

FindMarkers(microglia, ident.1 = "saline", ident.2 = "NMDA 48hr", group.by = "orig.ident", features = c("S1PR1", "ASAH1", "SGPL1", "SGPP1", "CERS5", "CERS6"), min.pct = 0.001, logfc.threshold = 0.001, test.use = "wilcox")

|++++++++++++++++++++++++++++++++++++++++++++++++++| 100% elapsed=00s

p_val	avg_log2FC pct.1 pct.2	p_val_adj

ASAH1 4.194271e-16 -0.726768391 0.154 0.510 7.817282e-12

SGPL1 2.912923e-11 -0.315981778 0.026 0.204 5.429106e-07

FindMarkers(microglia, ident.1 = "NMDA 12hr", ident.2 = "NMDA 48hr", group.by = "orig.ident", features = c("S1PR1", "ASAH1", "SGPL1", "SGPP1", "CERS5", "CERS6"), min.pct = 0.001, logfc.threshold = 0.001, test.use = "wilcox")

|++++++++++++++++++++++++++++++++++++++++++++++++++| 100% elapsed=00s

p_val avg_log2FC pct.1 pct.2	p_val_adj

ASAH1 1.892486e-08 -0.37066171 0.249 0.510 0.0003527216

**Cone Photoreceptors**

FindMarkers(cone_PRs, ident.1 = "saline", ident.2 = "NMDA 12hr", group.by = "orig.ident", features = c("S1PR1", "ASAH1", "SGPL1", "SGPP1", "CERS5", "CERS6"), min.pct = 0.001, logfc.threshold = 0.001, test.use = "wilcox")

|++++++++++++++++++++++++++++++++++++++++++++++++++| 100% elapsed=00s

p_val	avg_log2FC pct.1 pct.2	p_val_adj

ASAH1 1.757779e-23 0.750228175 0.582 0.472 3.276148e-19

FindMarkers(cone_PRs, ident.1 = "NMDA 3hr", ident.2 = "NMDA 12hr", group.by = "orig.ident", features = c("S1PR1", "ASAH1", "SGPL1", "SGPP1", "CERS5", "CERS6"), min.pct = 0.001, logfc.threshold = 0.001, test.use = "wilcox")

|++++++++++++++++++++++++++++++++++++++++++++++++++| 100% elapsed=00s

p_val	avg_log2FC pct.1 pct.2	p_val_adj

ASAH1 9.414644e-16 0.860505263 0.537 0.472 1.754701e-11

FindMarkers(cone_PRs, ident.1 = "NMDA 12hr", ident.2 = "NMDA 48hr", group.by = "orig.ident", features = c("S1PR1", "ASAH1", "SGPL1", "SGPP1", "CERS5", "CERS6"), min.pct = 0.001, logfc.threshold = 0.001, test.use = "wilcox")

|++++++++++++++++++++++++++++++++++++++++++++++++++| 100% elapsed=00s

p_val	avg_log2FC pct.1 pct.2	p_val_adj

ASAH1 3.046552e-29 -0.735056320 0.472 0.603 5.678164e-25

**Rod photoreceptors – nothing significant**

**Bipolar cells**

FindMarkers(BPs, ident.1 = "saline", ident.2 = "NMDA 3hr", group.by = "orig.ident", features = c("S1PR1", "ASAH1", "SGPL1", "SGPP1", "CERS5", "CERS6"), min.pct = 0.001, logfc.threshold = 0.001, test.use = "wilcox")

|++++++++++++++++++++++++++++++++++++++++++++++++++| 100% elapsed=00s

p_val avg_log2FC pct.1 pct.2	p_val_adj

S1PR1 1.112162e-09 0.16096240 0.026 0.006 2.072847e-05

FindMarkers(BPs, ident.1 = "saline", ident.2 = "NMDA 12hr", group.by = "orig.ident", features = c("S1PR1", "ASAH1", "SGPL1", "SGPP1", "CERS5", "CERS6"), min.pct = 0.001, logfc.threshold = 0.001, test.use = "wilcox")

|++++++++++++++++++++++++++++++++++++++++++++++++++| 100% elapsed=00s

p_val	avg_log2FC pct.1 pct.2	p_val_adj

S1PR1 4.041005e-17 0.185231193 0.026 0.005 7.531625e-13

CERS6 5.605735e-13 0.172260940 0.086 0.047 1.044797e-08

FindMarkers(BPs, ident.1 = "saline", ident.2 = "NMDA 48hr", group.by = "orig.ident", features = c("S1PR1", "ASAH1", "SGPL1", "SGPP1", "CERS5", "CERS6"), min.pct = 0.001, logfc.threshold = 0.001, test.use = "wilcox")

|++++++++++++++++++++++++++++++++++++++++++++++++++| 100% elapsed=00s

p_val	avg_log2FC pct.1 pct.2	p_val_adj

CERS6 1.738208e-07 0.203101850 0.086 0.056 0.003239671

FindMarkers(BPs, ident.1 = "NMDA 48hr", ident.2 = "NMDA 12hr", group.by = "orig.ident", features = c("S1PR1", "ASAH1", "SGPL1", "SGPP1", "CERS5", "CERS6"), min.pct = 0.001, logfc.threshold = 0.001, test.use = "wilcox")

|++++++++++++++++++++++++++++++++++++++++++++++++++| 100% elapsed=00s

p_val	avg_log2FC pct.1 pct.2 p_val_adj

S1PR1 7.416086e-07 0.068934884 0.015 0.005 0.0138221

**Horizontal cells – nothing significant**

**Amacrine cells**

FindMarkers(ACs, ident.1 = "saline", ident.2 = "NMDA 12hr", group.by = "orig.ident", features = c("S1PR1", "ASAH1", "SGPL1", "SGPP1", "CERS5", "CERS6"), min.pct = 0.001, logfc.threshold = 0.001, test.use = "wilcox")

|++++++++++++++++++++++++++++++++++++++++++++++++++| 100% elapsed=00s

p_val avg_log2FC pct.1 pct.2	p_val_adj

S1PR1 1.662805e-14 0.12475219 0.057 0.010 3.099137e-10

ASAH1 5.741345e-11 0.14428040 0.192 0.108 1.070072e-06

CERS6 9.634720e-09 0.10949798 0.136 0.074 1.795719e-04

SGPP1 4.425937e-05 0.08966504 0.062 0.033 8.249061e-01

**Retinal Ganglion cells**

FindMarkers(RGCs, ident.1 = "saline", ident.2 = "NMDA 3hr", group.by = "orig.ident", features = c("S1PR1", "ASAH1", "SGPL1", "SGPP1", "CERS5", "CERS6"), min.pct = 0.001, logfc.threshold = 0.001, test.use = "wilcox")

|++++++++++++++++++++++++++++++++++++++++++++++++++| 100% elapsed=00s

p_val avg_log2FC pct.1 pct.2	p_val_adj

ASAH1 1.662682e-08 0.30581668 0.676 0.460 0.0003098907

FindMarkers(RGCs, ident.1 = "saline", ident.2 = "NMDA 12hr", group.by = "orig.ident", features = c("S1PR1", "ASAH1", "SGPL1", "SGPP1", "CERS5", "CERS6"), min.pct = 0.001, logfc.threshold = 0.001, test.use = "wilcox")

|++++++++++++++++++++++++++++++++++++++++++++++++++| 100% elapsed=00s

p_val	avg_log2FC pct.1 pct.2	p_val_adj

CERS6 2.639325e-31 0.537813774 0.749 0.382 4.919173e-27

S1PR1 3.682310e-09 0.125082760 0.201 0.074 6.863089e-05

FindMarkers(RGCs, ident.1 = "saline", ident.2 = "NMDA 48hr", group.by = "orig.ident", features = c("S1PR1", "ASAH1", "SGPL1", "SGPP1", "CERS5", "CERS6"), min.pct = 0.001, logfc.threshold = 0.001, test.use = "wilcox")

|++++++++++++++++++++++++++++++++++++++++++++++++++| 100% elapsed=00s

p_val	avg_log2FC pct.1 pct.2	p_val_adj

CERS6 7.741105e-10 0.312013450 0.749 0.551 1.442787e-05

FindMarkers(RGCs, ident.1 = "NMDA 3hr", ident.2 = "NMDA 12hr", group.by = "orig.ident", features = c("S1PR1", "ASAH1", "SGPL1", "SGPP1", "CERS5", "CERS6"), min.pct = 0.001, logfc.threshold = 0.001, test.use = "wilcox")

|++++++++++++++++++++++++++++++++++++++++++++++++++| 100% elapsed=00s

p_val	avg_log2FC pct.1 pct.2	p_val_adj

CERS6 8.615019e-17 0.463978456 0.662 0.382 1.605667e-12

SGPL1 3.491665e-09 0.230269355 0.435 0.244 6.507764e-05

FindMarkers(RGCs, ident.1 = "NMDA 12hr", ident.2 = "NMDA 48hr", group.by = "orig.ident", features = c("S1PR1", "ASAH1", "SGPL1", "SGPP1", "CERS5", "CERS6"), min.pct = 0.001, logfc.threshold = 0.001, test.use = "wilcox")

|++++++++++++++++++++++++++++++++++++++++++++++++++| 100% elapsed=00s

p_val	avg_log2FC pct.1 pct.2	p_val_adj

CERS6 1.578833e-09 -0.225800324 0.382 0.551 2.942629e-05
