## Supplemental Table 2 for "Sphingosine-1-phosphate signaling regulates the ability of Müller glia to become neurogenic, proliferating progenitor-like cells"

**Supplemental table – Statistics for dotplots**

**Figure 7j - stats Muller Glia clod-treated for S1P-related genes**

FindMarkers(MG, ident.1 = "SS", ident.2 = "clod_SS", group.by = "orig.ident", features = c("S1PR1", "SPHK1", "ASAH1", "SGPL1", "CERS5", "CERS6"), min.pct = 0.001, logfc.threshold = 0.001, test.use = "wilcox")

|++++++++++++++++++++++++++++++++++++++++++++++++++| 100% elapsed=00s

p_val	avg_log2FC pct.1 pct.2	p_val_adj

S1PR1 9.969211e-21 0.537367807 0.626 0.424 2.100114e-16

SPHK1 1.778776e-04 0.188524388 0.160 0.100 1.000000e+00

ASAH1 3.083628e-03 0.161901419 0.275 0.215 1.000000e+00

CERS6 1.460267e-01 -0.227103669 0.246 0.263 1.000000e+00

SGPL1 4.146942e-01 0.006208824 0.139 0.126 1.000000e+00

CERS5 4.432017e-01 0.016230049 0.067 0.058 1.000000e+00

FindMarkers(MG, ident.1 = "SS", ident.2 = "NM", group.by = "orig.ident", features = c("S1PR1", "SPHK1", "ASAH1", "SGPL1", "CERS5", "CERS6"), min.pct = 0.001, logfc.threshold = 0.001, test.use = "wilcox")

|++++++++++++++++++++++++++++++++++++++++++++++++++| 100% elapsed=00s

p_val avg_log2FC pct.1 pct.2	p_val_adj

S1PR1 8.779736e-137 1.74259721 0.626 0.161 1.849539e-132

CERS6 2.253022e-17 0.49540617 0.246 0.123 4.746215e-13

CERS5 1.160099e-11 -0.13590893 0.067 0.168 2.443865e-07

SPHK1 1.803931e-02 0.23242095 0.160 0.140 1.000000e+00

ASAH1 6.891110e-01 0.11600914 0.275 0.312 1.000000e+00

SGPL1 7.712523e-01 0.08387772 0.139 0.157 1.000000e+00

FindMarkers(MG, ident.1 = "NM", ident.2 = "clod_NM", group.by = "orig.ident", features = c("S1PR1", "SPHK1", "ASAH1", "SGPL1", "CERS5", "CERS6"), min.pct = 0.001, logfc.threshold = 0.001, test.use = "wilcox")

|++++++++++++++++++++++++++++++++++++++++++++++++++| 100% elapsed=00s

p_val avg_log2FC pct.1 pct.2	p_val_adj

S1PR1 1.879058e-12 -0.50158870 0.161 0.290 3.958423e-08

CERS5 4.708781e-06 0.12197213 0.168 0.083 9.919518e-02

CERS6 8.752308e-04 -0.23453203 0.123 0.176 1.000000e+00

SPHK1 3.004105e-01 -0.02699285 0.140 0.118 1.000000e+00

SGPL1 3.333511e-01 0.01698714 0.157 0.134 1.000000e+00

ASAH1 9.777293e-01 -0.07677070 0.312 0.289 1.000000e+00

FindMarkers(MG, ident.1 = "clod_SS", ident.2 = "clod_NM", group.by = "orig.ident", features = c("S1PR1", "SPHK1", "ASAH1", "SGPL1", "CERS5", "CERS6"), min.pct = 0.001, logfc.threshold = 0.001, test.use = "wilcox")

|++++++++++++++++++++++++++++++++++++++++++++++++++| 100% elapsed=00s

p_val avg_log2FC pct.1 pct.2	p_val_adj

S1PR1 2.322821e-12 0.70364070 0.424 0.290 4.893255e-08

CERS6 1.577835e-06 0.48797782 0.263 0.176 3.323866e-02

ASAH1 1.571366e-02 -0.12266298 0.215 0.289 1.000000e+00

CERS5 8.355572e-02 -0.03016685 0.058 0.083 1.000000e+00

SPHK1 4.269028e-01 0.01690371 0.100 0.118 1.000000e+00

SGPL1 9.377571e-01 0.09465604 0.126 0.134 1.000000e+00

**Figure 7k - stats amacrine cells clod-treated for S1P-related genes**

FindMarkers(ACs, ident.1 = "SS", ident.2 = "clod_SS", group.by = "orig.ident", features = c("S1PR1", "SPHK1", "ASAH1", "SGPL1", "CERS5", "CERS6"), min.pct = 0.001, logfc.threshold = 0.001, test.use = "wilcox")

|++++++++++++++++++++++++++++++++++++++++++++++++++| 100% elapsed=00s

p_val avg_log2FC pct.1 pct.2 p_val_adj

CERS5 0.0000266165 -0.09806320 0.072 0.106 0.5607033

CERS6 0.0030325528 -0.04632226 0.080 0.104 1.0000000

S1PR1 0.0426806443 0.09174008 0.039 0.029 1.0000000

SPHK1 0.4737358962 0.09610572 0.048 0.045 1.0000000

ASAH1 0.6778338914 0.12356340 0.092 0.091 1.0000000

SGPL1 0.7333057245 0.05117452 0.068 0.072 1.0000000

FindMarkers(ACs, ident.1 = "SS", ident.2 = "NM", group.by = "orig.ident", features = c("S1PR1", "SPHK1", "ASAH1", "SGPL1", "CERS5", "CERS6"), min.pct = 0.001, logfc.threshold = 0.001, test.use = "wilcox")

|++++++++++++++++++++++++++++++++++++++++++++++++++| 100% elapsed=00s

p_val avg_log2FC pct.1 pct.2	p_val_adj

SGPL1 1.084968e-07 -0.14599684 0.068 0.118 0.002285593

SPHK1 2.118928e-03 -0.02347995 0.048 0.072 1.000000000

ASAH1 3.076202e-03 -0.02824744 0.092 0.124 1.000000000

CERS5 2.384050e-02 -0.06902253 0.072 0.093 1.000000000

S1PR1 8.194360e-02 0.09171492 0.039 0.029 1.000000000

FindMarkers(ACs, ident.1 = "NM", ident.2 = "clod_NM", group.by = "orig.ident", features = c("S1PR1", "SPHK1", "ASAH1", "SGPL1", "CERS5", "CERS6"), min.pct = 0.001, logfc.threshold = 0.001, test.use = "wilcox")

|++++++++++++++++++++++++++++++++++++++++++++++++++| 100% elapsed=00s

p_val avg_log2FC pct.1 pct.2 p_val_adj

ASAH1 6.810306e-07 0.31915078 0.124 0.075 0.01434659

SGPL1 2.083544e-02 0.16155783 0.118 0.095 1.00000000

S1PR1 1.365262e-01 -0.04394308 0.029 0.038 1.00000000

SPHK1 4.056890e-01 0.03815822 0.072 0.065 1.00000000

CERS6 6.578706e-01 0.04301406 0.090 0.096 1.00000000

CERS5 7.498299e-01 0.11272050 0.093 0.091 1.00000000

FindMarkers(ACs, ident.1 = "clod_SS", ident.2 = "clod_NM", group.by = "orig.ident", features = c("S1PR1", "SPHK1", "ASAH1", "SGPL1", "CERS5", "CERS6"), min.pct = 0.001, logfc.threshold = 0.001, test.use = "wilcox")

|++++++++++++++++++++++++++++++++++++++++++++++++++| 100% elapsed=00s

p_val avg_log2FC pct.1 pct.2 p_val_adj

SPHK1 0.002166248 -0.08142745 0.045 0.065	1

SGPL1 0.005420232 -0.03561354 0.072 0.095	1

ASAH1 0.033009855 0.16733993 0.091 0.075	1

CERS5 0.056829516 0.14176117 0.106 0.091	1

S1PR1 0.094143064 -0.04396824 0.029 0.038	1

CERS6 0.304055259 0.09021781 0.104 0.096	1

**Figure 7l - stats bipolar cells clod-treated for S1P-related genes**

**Nothing significant**

**Figure 7m - stats retinal ganglion cells clod-treated for S1P-related genes**

FindMarkers(RGCs, ident.1 = "NM", ident.2 = "clod_NM", group.by = "orig.ident", features = c("S1PR1", "SPHK1", "ASAH1", "SGPL1", "CERS5", "CERS6"), min.pct = 0.001, logfc.threshold = 0.001, test.use = "wilcox")

|++++++++++++++++++++++++++++++++++++++++++++++++++| 100% elapsed=00s

p_val avg_log2FC pct.1 pct.2	p_val_adj

ASAH1 2.209078e-08 0.24561769 0.479 0.317 0.0004653644

SPHK1 3.096344e-07 0.12849418 0.190 0.088 0.0065227574

CERS6 1.698275e-03 -0.15830403 0.441 0.499 1.0000000000

S1PR1 1.106918e-02 0.06874279 0.202 0.141 1.0000000000

CERS5 4.695818e-02 0.04236618 0.146 0.106 1.0000000000

SGPL1 1.061835e-01 0.03188283 0.171 0.132 1.0000000000
