## Supplementary figures and images for "Sphingosine-1-phosphate signaling regulates the ability of Müller glia to become neurogenic, proliferating progenitor-like cells"

### Supplemental Fig. 1

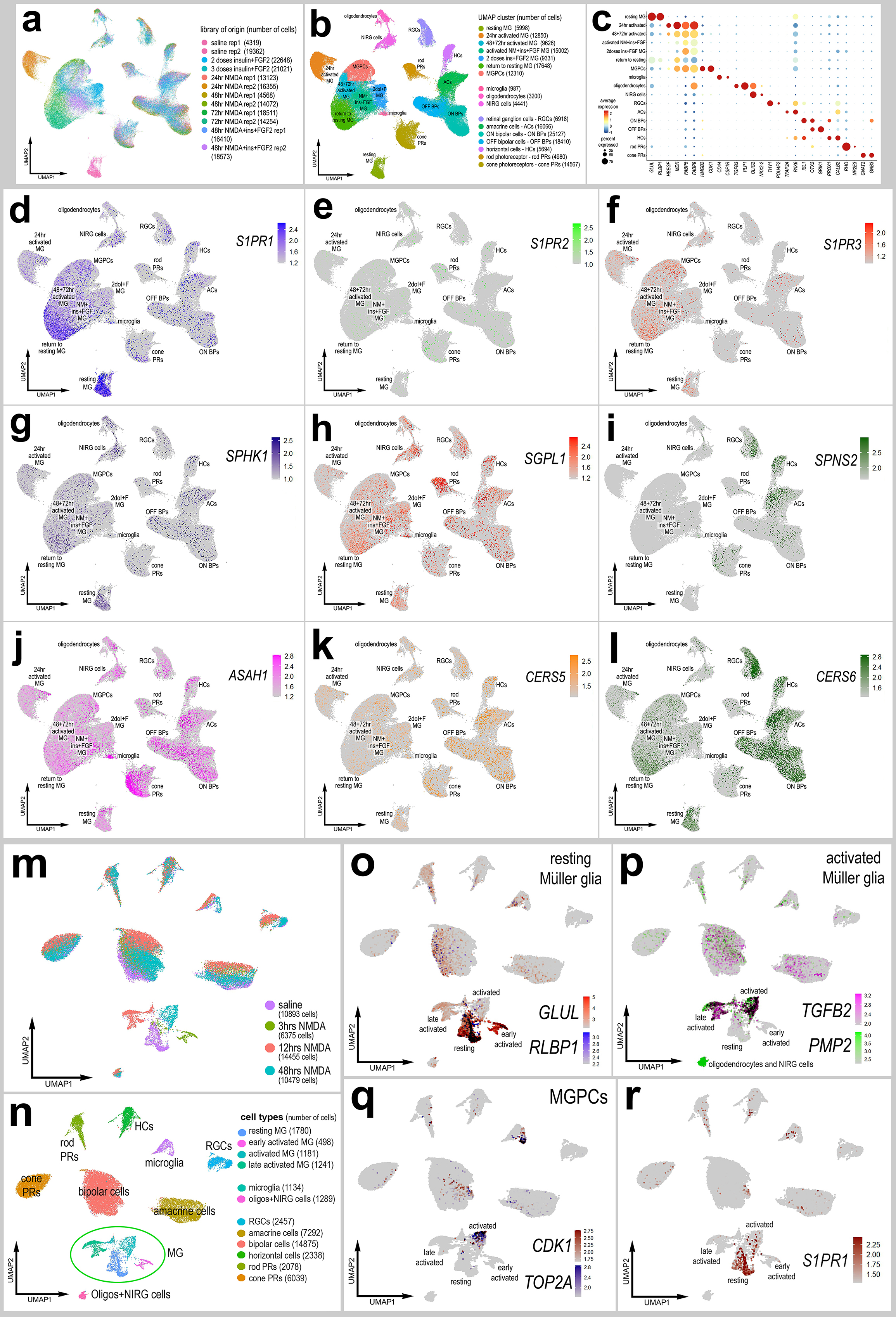

### Supplemental Fig. 2

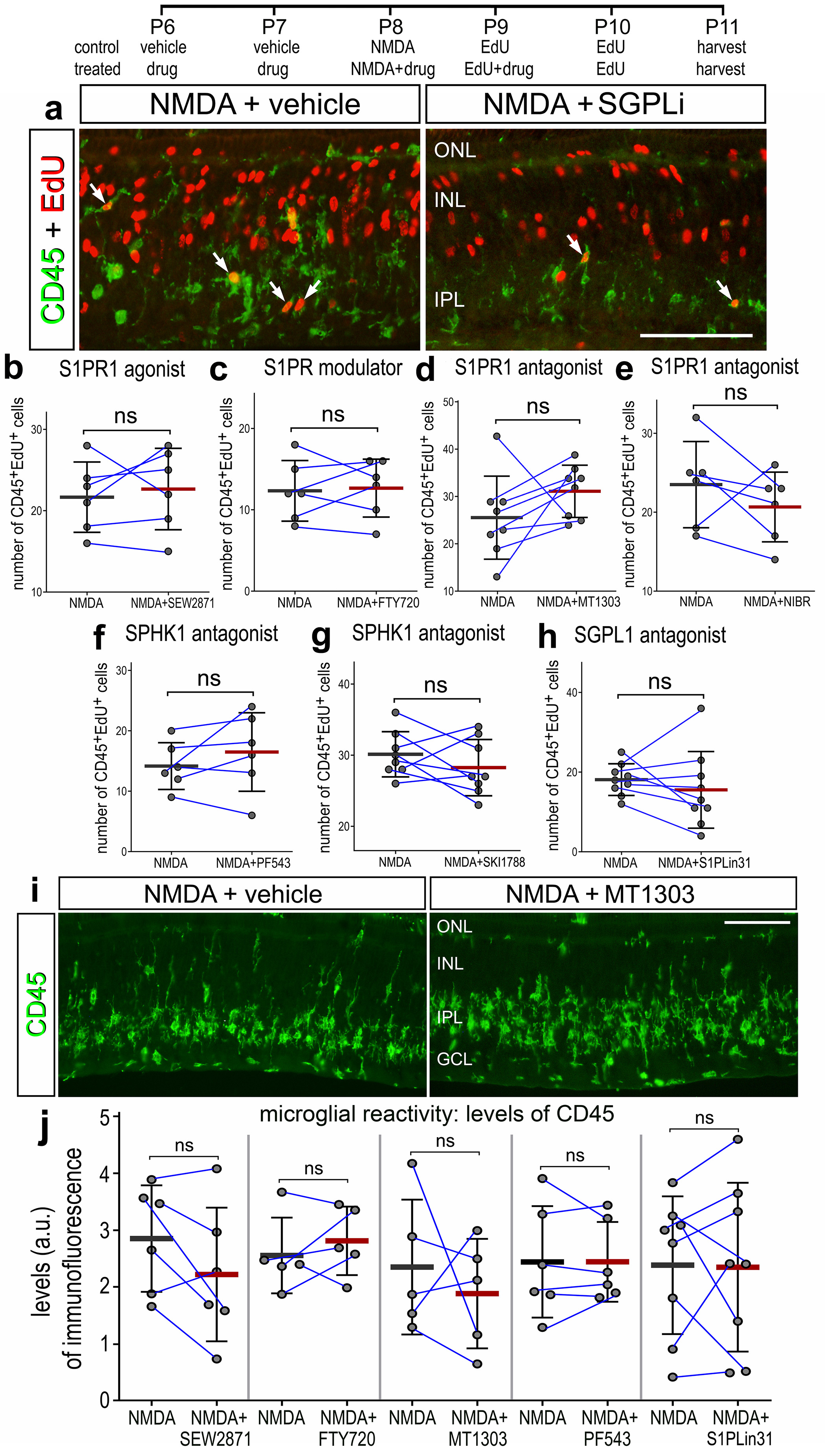

### Supplemental Fig. 3

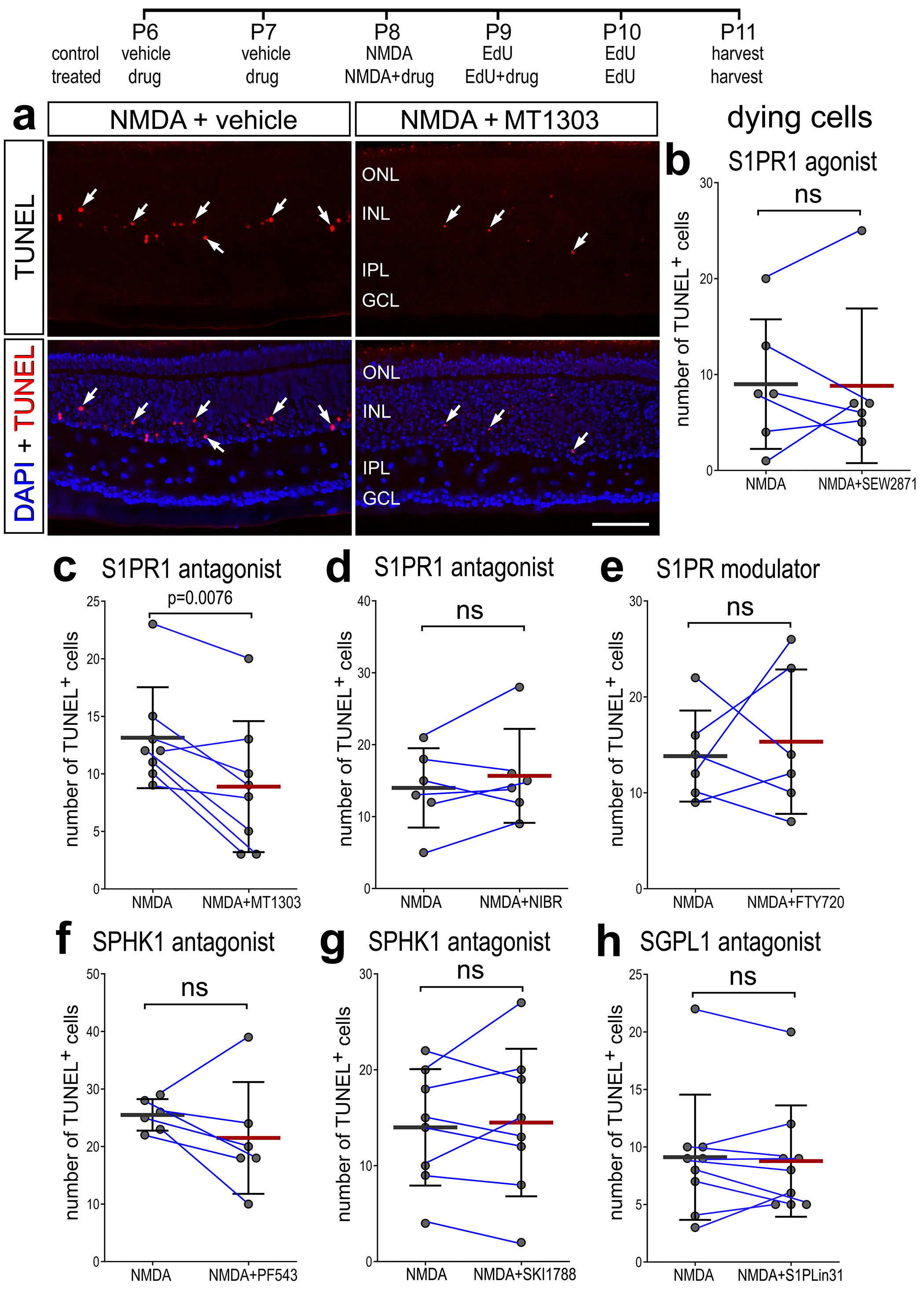

### Supplemental Fig. 4

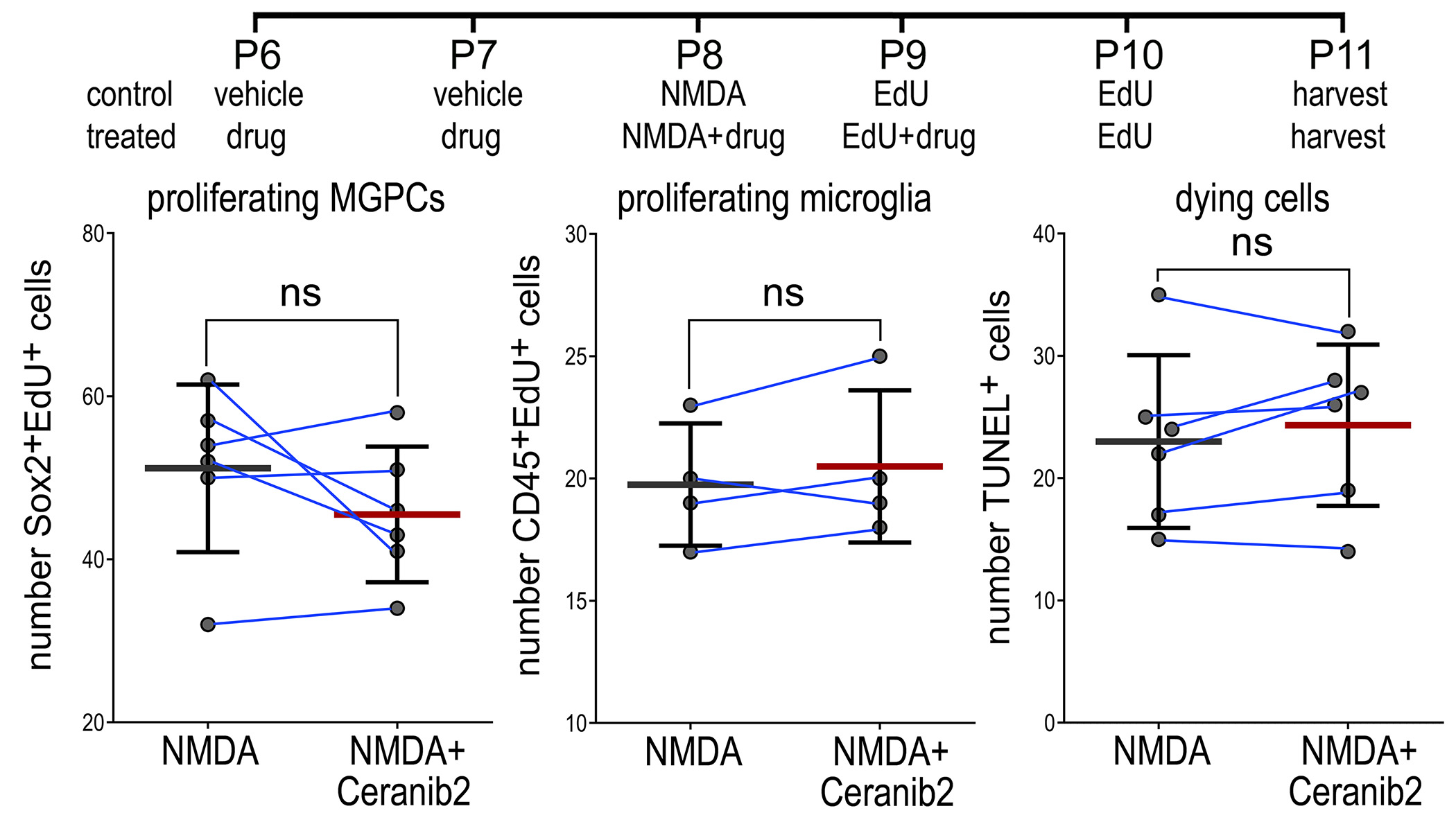

### Supplemental Fig. 5

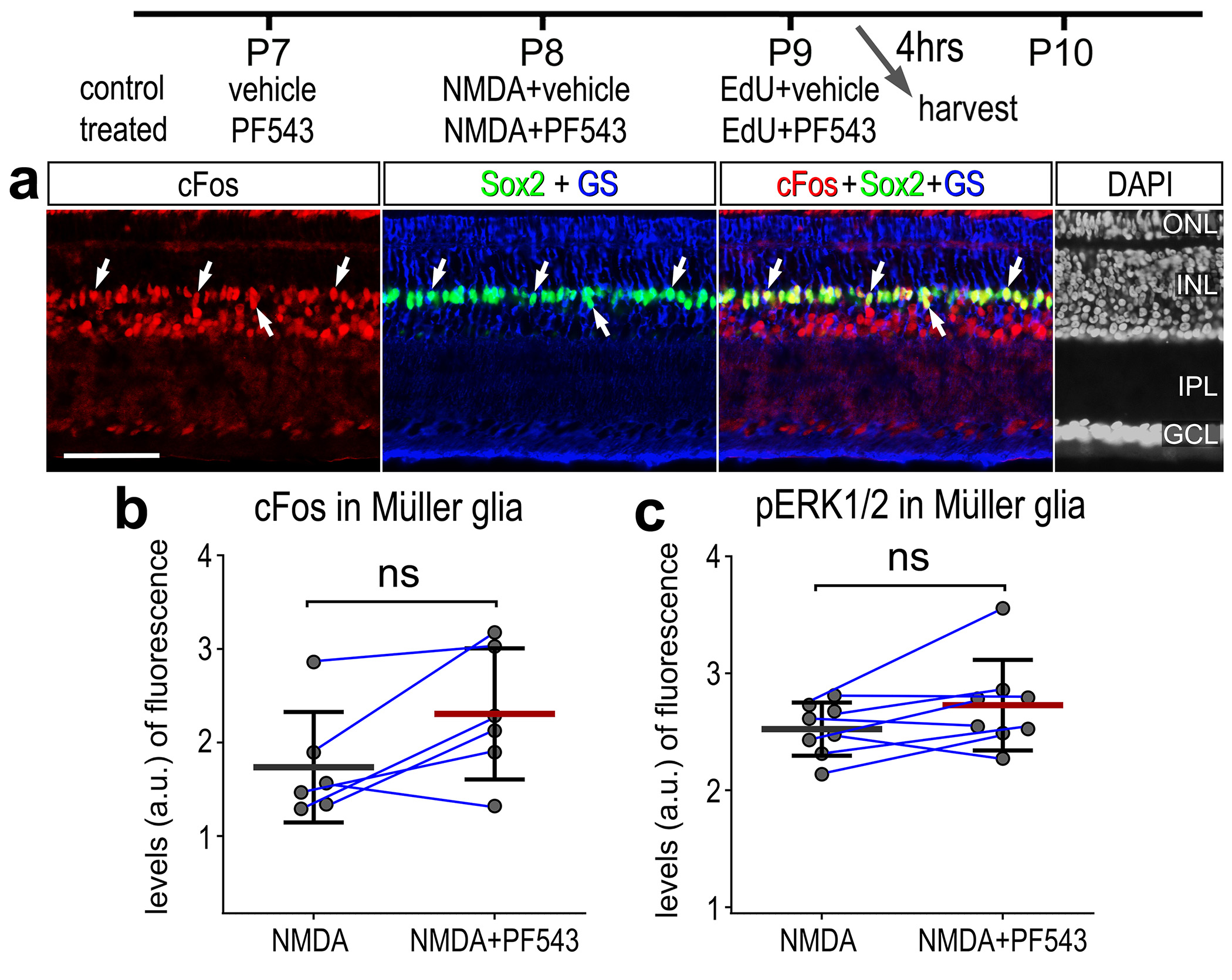

### Supplemental Fig. 6

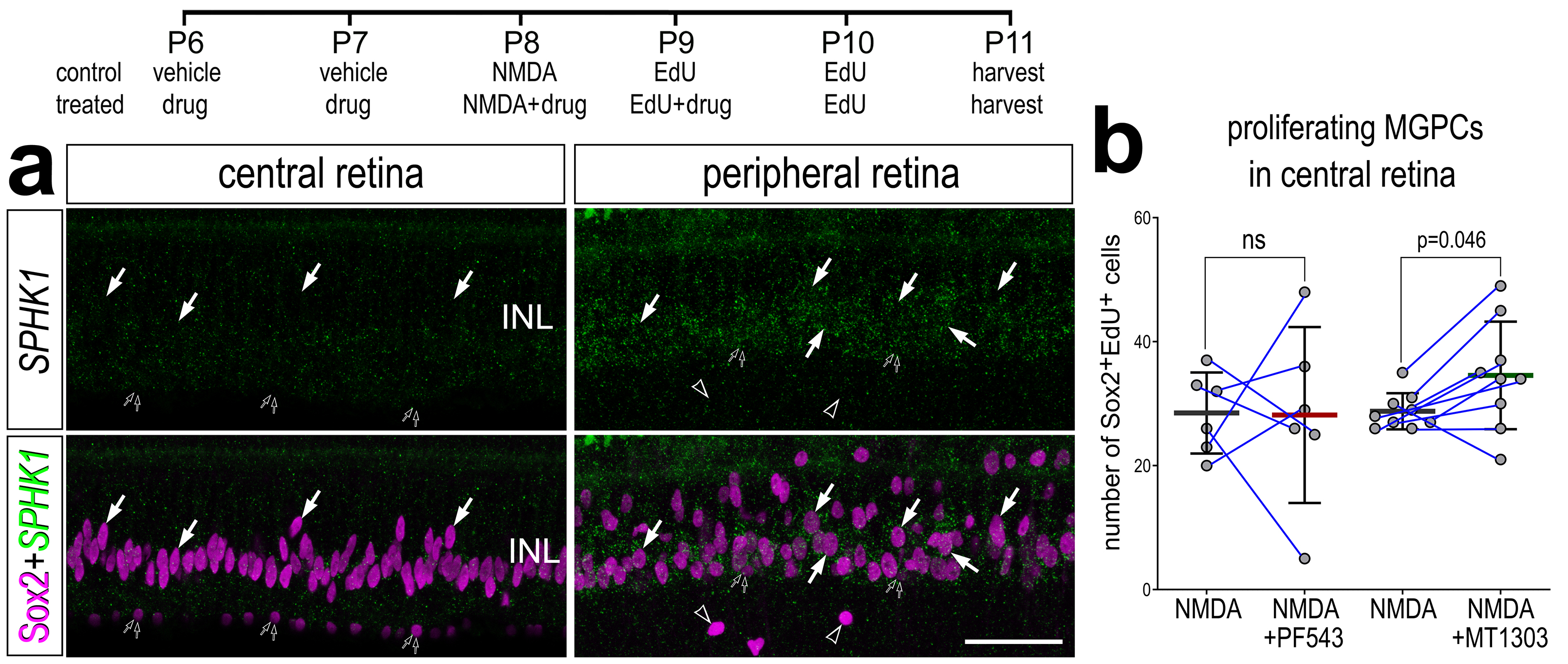
